# Multiple QTL underlie milk phenotypes at the CSF2RB locus

**DOI:** 10.1101/414821

**Authors:** Thomas J Lopdell, Kathryn Tiplady, Christine Couldrey, Thomas JJ Johnson, Michael Keehan, Stephen R Davis, Bevin L Harris, Richard J Spelman, Russell G Snell, Mathew D Littlejohn

## Abstract

**Background:** Bovine milk provides an important source of nutrition in much of the Western world, forming components of many food products. Over many years, artificial selection has substantially improved milk production by cows. However, the genes underlying milk production quantitative trait loci (QTL) remain relatively poorly characterised. Here, we investigate a previously-reported QTL located at the *CSF2RB* locus, for several milk production phenotypes, to better understand its underlying genetic and molecular causes.

**Results:** Using a population of 29,350 taurine dairy cattle, we conducted association analyses for milk yield and composition traits, and identified highly significant QTL for milk yield, milk fat concentration, and milk protein concentration. Strikingly, protein concentration and milk yield appear to show co-located yet genetically distinct QTL. To attempt to understand the molecular mechanisms that might be mediating these effects, gene expression data were used to investigate eQTL for eleven genes in the broader interval. This analysis highlighted genetic impacts on *CSF2RB* and *NCF4* expression that share similar association signatures to those observed for lactation QTL, strongly implicating one or both of these genes as the cause of these effects. Using the same gene expression dataset representing 357 lactating cows, we also identified 38 novel RNA editing sites in the 3^*′*^ UTR of *CSF2RB* transcripts. The extent to which two of these sites were edited also appears to be genetically co-regulated with lactation QTL, highlighting a further layer of regulatory complexity implicating the *CSF2RB* gene.

**Conclusions:** This chromosome 5 locus presents a diversity of molecular and lactation QTL, likely representing multiple overlapping effects that, at a minimum, highlight the *CSF2RB* gene as having a causal role in these processes.

## Background

Liquid milk provides a source of nutrition to neonate mammals, and is also used as a convenient source of nutrition for both infant and adult humans. In much of the Western world, milk is primarily produced for human consumption by taurine cattle (*Bos taurus*) dairy breeds. Within these breeds, many generations of selection have improved milk production capacity and efficiency. However, despite numerous recent genome-wide association studies (GWAS) e.g. [1, 2, 3, 4], major QTL remain for which no causative gene has been definitively assigned.

Several genes with substantial impacts on milk yield are known, including *DGAT1* [5], *ABCG2* [6], *GHR* [7], *SLC37A1* [8], and *MGST1* [9]. Recently, as part of work presented elsewhere [10], we performed a milk volume genome-wide association analysis in 4,982 mixed breed cattle using a BayesB model [11, 12], using a panel of 3,695 variants selected as tag-SNPs representing expression QTL (eQTL) from lactating mammary tissue. Of the top three loci explaining the greatest proportion of genetic variance in this model, genes representing the top and second to top effects have been well described for their role in milk production effects (*DGAT1* and *MGST1* respectively [5, 9]), whereas no causative gene appears to have been definitively assigned for the third signal at chromosome 5:75–76 Mbp.

This locus broadly overlaps QTL reported previously for milk yield [13, 3], milk protein yield [13, 3], milk protein concentration [14, 1, 2], and milk fat concentration [9, 2].Although no gene has been definitively implicated, Pausch et al [2] noted significant markers mapping adjacent to the *CSF2RB, NCF4* genes, and *TST* genes, proposing the latter as the most likely candidate based on its proximity to the top associated variant. Other studies have proposed *CSF2RB* due to its high level of expression in the mammary gland [14, 1], or involvement in the JAK-STAT signalling pathway [13, 3]. Other nearby genes speculated to cause these effects also include *MYH9* [3] and *NCF4* [13].

Given these observations, and the magnitude and diversity of effects at this locus, the aim of this study was to investigate the chromosome 5 region in detail. By combining information from milk yield and composition traits with gene expression data from a large bovine mammary RNA sequence dataset, we highlight multiple lactation, gene expression, and RNA-editing QTL segregating at the locus, presenting *CSF2RB* as the most likely causative gene responsible for these effects.

## Results

### Sequence-based association analysis at the chr5 interval

Fine mapping of milk yield and protein concentration QTL at the chr5:75–76Mbp locus was performed using imputed sequence genotypes representing >30,000 cows. These animals had been physically genotyped using the GeneSeek Genomic Profiler (GGP) chip, where this panel had also been augmented with 224 custom sequence variants representing the chromosome 5 window, enriching the interval for QTL-tag variants identified from previous, preliminary analyses of the locus (spanning 74.8–76.2Mbp; see Materials and Methods). Sequence data were imputed using Beagle4 [15] (74.8–76.2Mbp; 11,733 markers), and phenotypes were produced from herd-test records (N=29,350 cows) from the animals’ first lactations to derive values for milk yield (MY), protein yield (PY), fat yield (FY), protein concentration (PC), and fat concentration (FC; see Materials and Methods).

Mixed linear model association (MLMA) analyses were conducted using GCTA [16]. The top associated variant for each of the five phenotypes is shown in Table 1. All QTL were found to be significant at the genome-wide threshold 5 × 10^−8^. The most significant QTL was identified for protein concentration, followed by fat concentration and milk yield. The fat yield phenotype exhibited the least significant QTL. The protein concentration and milk yield phenotype QTL are illustrated in Figure 1.

**Table 1.** Top variants for milk yield and composition trait QTL. Phenotypes are daily yields for fat (FY), protein (PY), and milk (MY); and composition (percentage) phenotypes for fat (FC) and protein (PC).

| Phenotype | Top Variant | MAF | Beta | SE | P |
| --- | --- | --- | --- | --- | --- |
| FY (kg/day) | rs466308089 | 0.031 | -0.015 | 0.003 | $2.40 \times 10^{-8}$ |
| PY (kg/day) | rs108985709 | 0.409 | 0.004 | 0.001 | $1.05 \times 10^{-8}$ |
| MY (L/day) | rs208473130 | 0.444 | 0.216 | 0.021 | $6.64 \times 10^{-25}$ |
| FC (%) | rs379739117 | 0.471 | -0.055 | 0.004 | $3.27 \times 10^{-41}$ |
| PC (%) | rs208375076 | 0.446 | -0.035 | 0.002 | $7.28 \times 10^{-83}$ |

**Figure 1.**
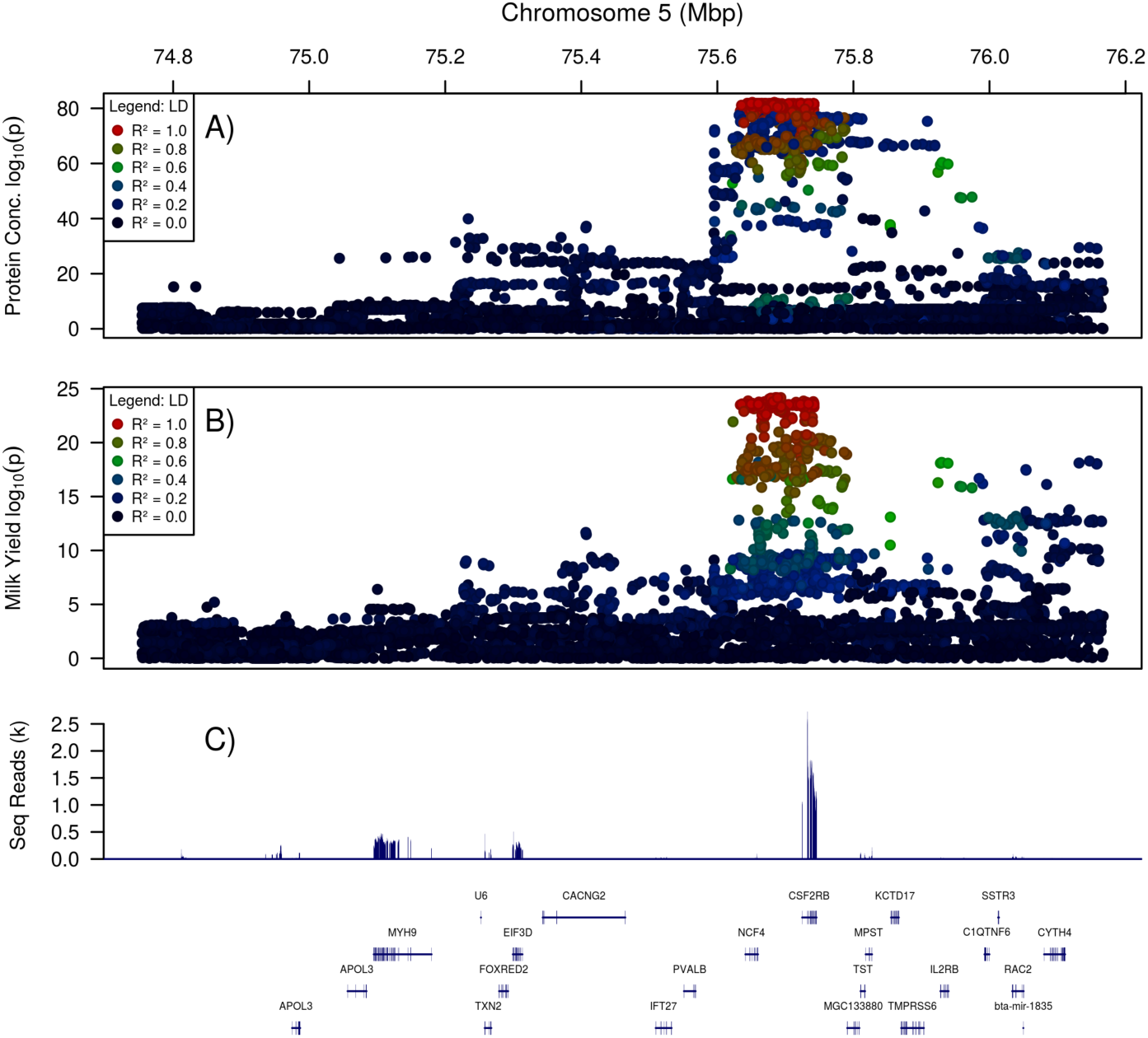
The genetic context of milk trait QTLs. Panels A) and B): QTLs for the herd-test-derived phenotypes protein concentration (A) and milk yield (B). Colours represent LD (*R*^2^) with the most significant marker. Panel C) shows the locations of genes mapping into this window (bottom) and the numbers of RNAseq reads mapping at positions across the window (top).

Alongside the MLMA-LOCO analysis, AI-REML was performed, using a GRM calculated over all chromosomes, to estimate narrow-sense heritabilities (*h*^2^; Table 2). To investigate these QTL further, the linkage disequilibrium (LD) statistics (*R*^2^) between each pair of top variants were calculated (Figure 2). Strong LD was observed between the top variants for the MY, FC, and PC phenotypes (MY vs FC tag variants *R*^2^ = 0.887; MY vs PC tag variants *R*^2^ = 0.991).

**Table 2.** Heritability estimates for milk yield and composition phenotypes. Phenotypes are milk fat daily yield (kg) and concentration (%), protein daily yield and concentration, and milk daily volume (L).

| FY (kg) | FC (%) | PY (kg) | PC (%) | MY (L) |
| --- | --- | --- | --- | --- |
| 0.184 ± 0.008 | 0.622 ± 0.007 | 0.183 ± 0.008 | 0.614 ± 0.007 | 0.263 ± 0.008 |

**Figure 2.**
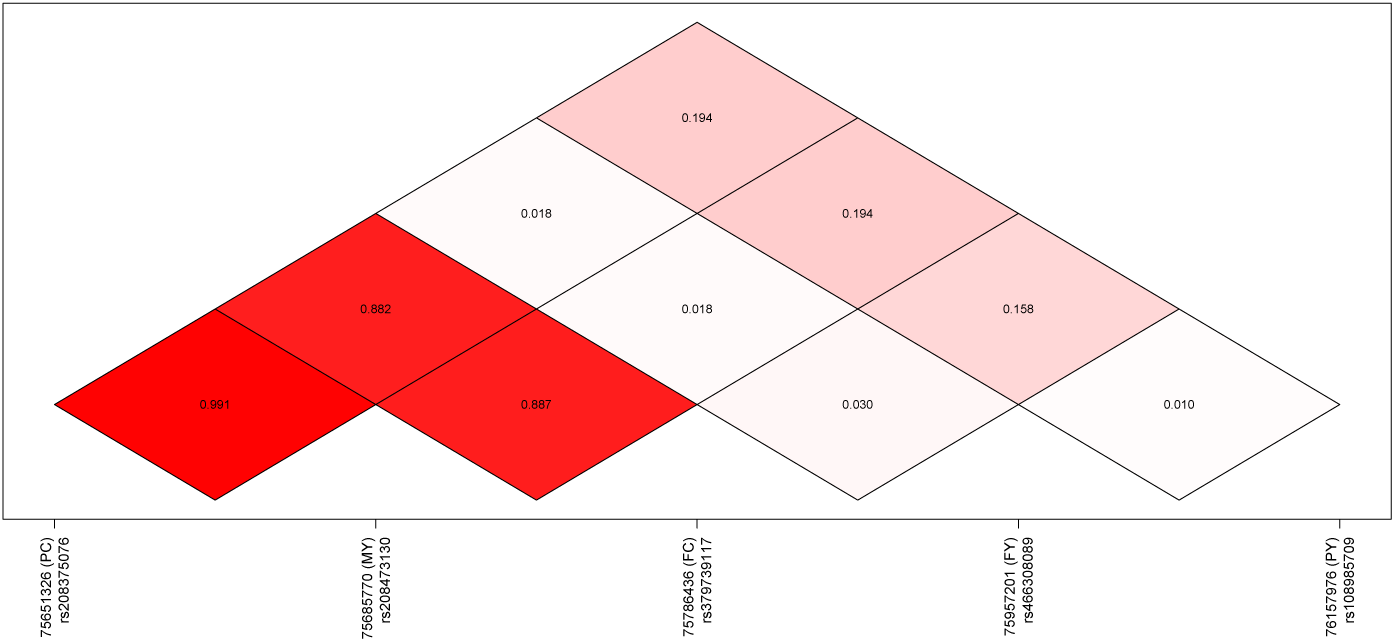
Linkage disequilibrium (LD) observed between the top associated markers for each phenotype (*R*^*2*^). Markers are identified using dbSNP reference SNP ID numbers. Phenotypes are as per Table 2.

### Functional prediction of variant effects suggests regulatory QTL mechanisms

To assess potential functional effects of the statistically implicated QTL variants, all polymorphisms in strong LD (*R*^2^ *>* 0.9) with the top-ranked QTL variants for each trait were extracted (N=365 variants), and analysed using the Ensembl Variant Effect Predictor (VEP) [17]. The majority of these variants (N=247) were predicted to map outside of genes, while 113 were predicted to be intronic, with 58 in transcript ENSBTAT00000009911.4 (*NCF4*) and 55 in ENSBTAT00000011947.5 (*CSF2RB*).The remaining five variants were predicted to be synonymous mutations, with two in ENSBTAP00000009911.4 (*NCF4*) at positions p.Gln145= and p.Tyr243=, and three in ENSBTAP00000011947.5 (*CSF2RB*) at positions p.Asn58=, p.Tyr405=, and p.Glu424=. Importantly, none of the highly associated variants were predicted to change the protein sequences of genes, suggesting a regulatory mode of effect as the likely mechanism(s) of the QTL.

### Expression QTL analysis highlights three genes differentially expressed by genotype

To look for *cis*-eQTL effects that might explain the lactation QTL, gene expression levels were calculated for genes in the BTA5:75-76Mbp window, using RNAseq data representing lactating mammary tissue biopsies from 357 cows (Figure 1, panel C). Expression levels in fragments per kilobase of transcript per million mapped reads (FPKM) and transcripts per million mapped reads (TPM) were calculated using Stringtie [18] and are shown in Table 3 for transcripts where FPKM *>* 0.1. The gene with the highest expression level was *CSF2RB*, consistent with previous observations in murine mammary RNAseq data [19]. Moderate expression was also observed for the candidate gene *MYH9*. However, the expression level of *NCF4* was very low, at FPKM = 0.406. The highest correlation between pairs of gene expression levels was observed for *TST* and *MPST* (*r* = 0.545), concordant with the published observation of a shared bidirectional promoter for these two genes [20].

**Table 3.**
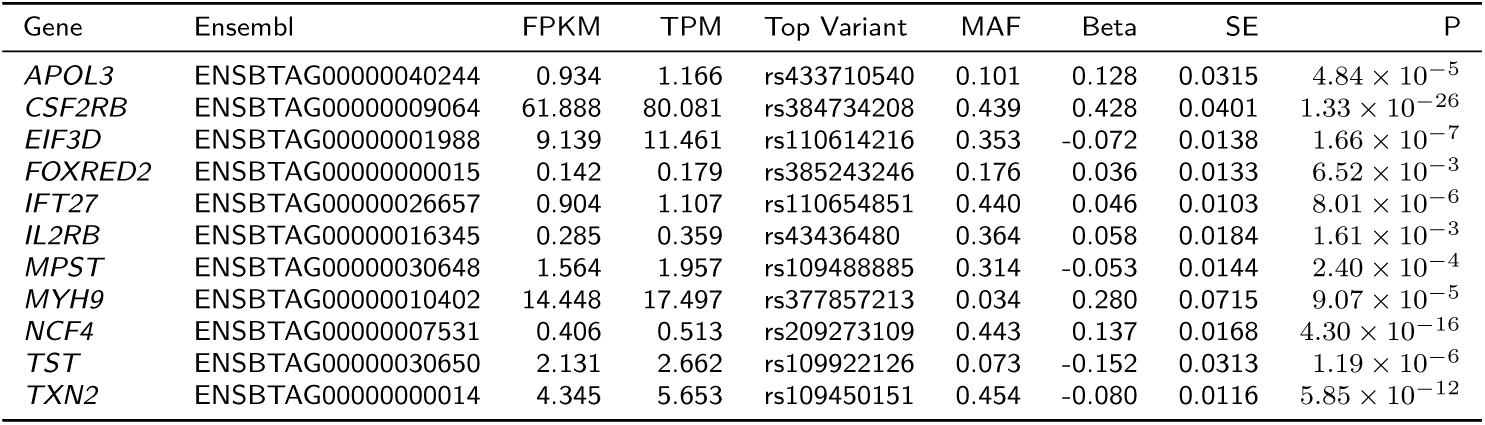
Median gene expression levels and top variants identified in eQTL analyses. Genes with FPKM values less than 0.1 are not shown. Gene symbols are from VGNC and Ensembl. Beta is the effect size of the minor allele on gene expression, measured in VST-transformed units. Three genes have eQTL which exceed the genome-wide significance threshold 5 × 10^−8^ [60].

Association mapping was then conducted for the eleven expressed genes in Table 3. To this end, gene expression data were first scaled using the variance-stabilising transformation (VST) implemented in DESeq [21]. A GRM was then calculated for the 357 cows representing the RNAseq dataset, and the MLMA-LOCO method performed as described for analysis of lactation traits, above. This yielded genome-wide significant eQTL for three genes: *CSF2RB* (1.33 × 10^−26^), *NCF4* (4.30 × 10^−16^), and *TXN2* (5.85 × 10^−12^) (see Table 3 and Figure 3). All three genes locate within the peaks of their respective eQTL, demonstrating regulation in *cis*.

**Figure 3.**
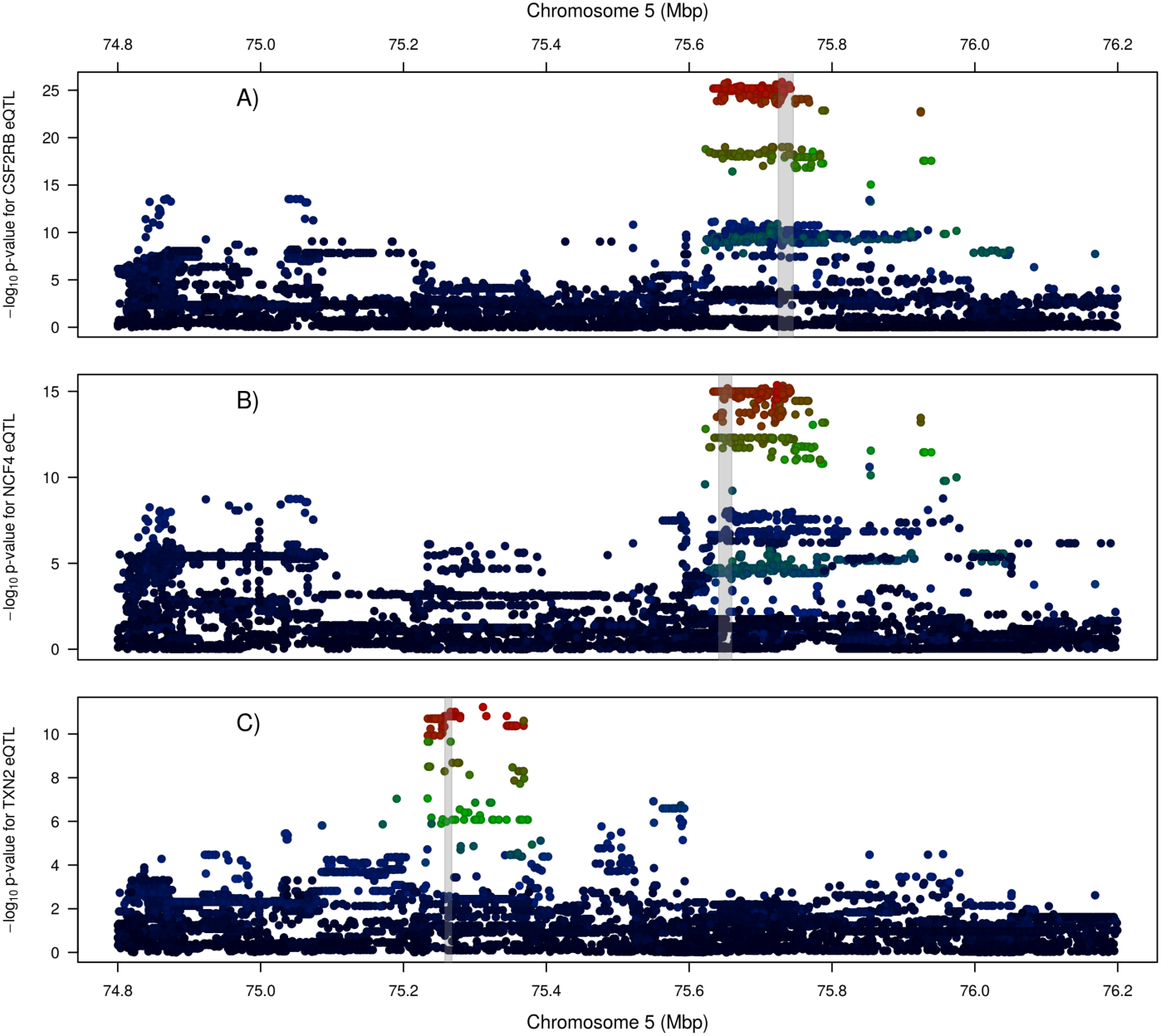
QTL plots showing eQTL for the three genes that exhibit genome-wide significant *cis*-eQTL (Table 3). From top to bottom, the three genes are A) *CSF2RB*, B) *NCF4*, and C) *TXN2*. Colours represent correlations for each marker with the top variant for that eQTL (see Figure 1 for legend). Grey bands indicate the location of the gene for which the eQTL is displayed.

In cases where genetic regulation of gene expression (i.e. an eQTL) underlies a complex trait QTL, we expect that both QTL should share similar association signals, with the most (and least) associated variants similar between phenotypes. To test whether any of the eleven expressed genes shared similarities with the milk QTL, the Pearson correlation between the log_10_ p-values for each of the milk QTL and eQTL were calculated. Table 4 shows the QTL:eQTL correlations for all five phenotypes with three significant eQTL, plus the *TST* gene, which did not yield a genome-wide significant eQTL, though has been proposed as a candidate underlying this locus. The eQTL for *CSF2RB* has *R*^2^ *>* 0.5 (*r >* 0.707) with three of the five milk phenotypes, while correlations for the neighbouring gene *NCF4* sit just below this level. Neither of the *TXN2* or *TST* genes exhibited high correlations with any milk QTL. The eQTL for *CSF2RB* was also highly correlated with the *NCF4* eQTL (*r* = 0.863). A similar picture is obtained when examining the LD between the top tag markers for each QTL, with high LD observed (Figure 4) among the tags for the MY, FC, and PC phenotypes with the tags for the *CSF2RB* and *NCF4* eQTL.

**Table 4.** Pearson correlations between the – log_10_ p-values for milk trait QTL and co-located gene expression QTL. Three genes with significant (P<5 10^−8^) eQTL are shown, along with the *TST* gene [2] that have previously been proposed as a candidate causative at this locus.

| Phenotype | *CSF2RB* | *NCF4* | *TST* | *TXN2* |
| --- | --- | --- | --- | --- |
| FY (kg/day) | 0.376 | 0.164 | 0.293 | 0.024 |
| PY (kg/day) | 0.562 | 0.404 | 0.425 | 0.032 |
| MY (L/day) | 0.849 | 0.682 | 0.306 | -0.039 |
| FC (%) | 0.756 | 0.648 | 0.293 | -0.128 |
| PC (%) | 0.754 | 0.689 | 0.059 | -0.118 |

**Figure 4.**
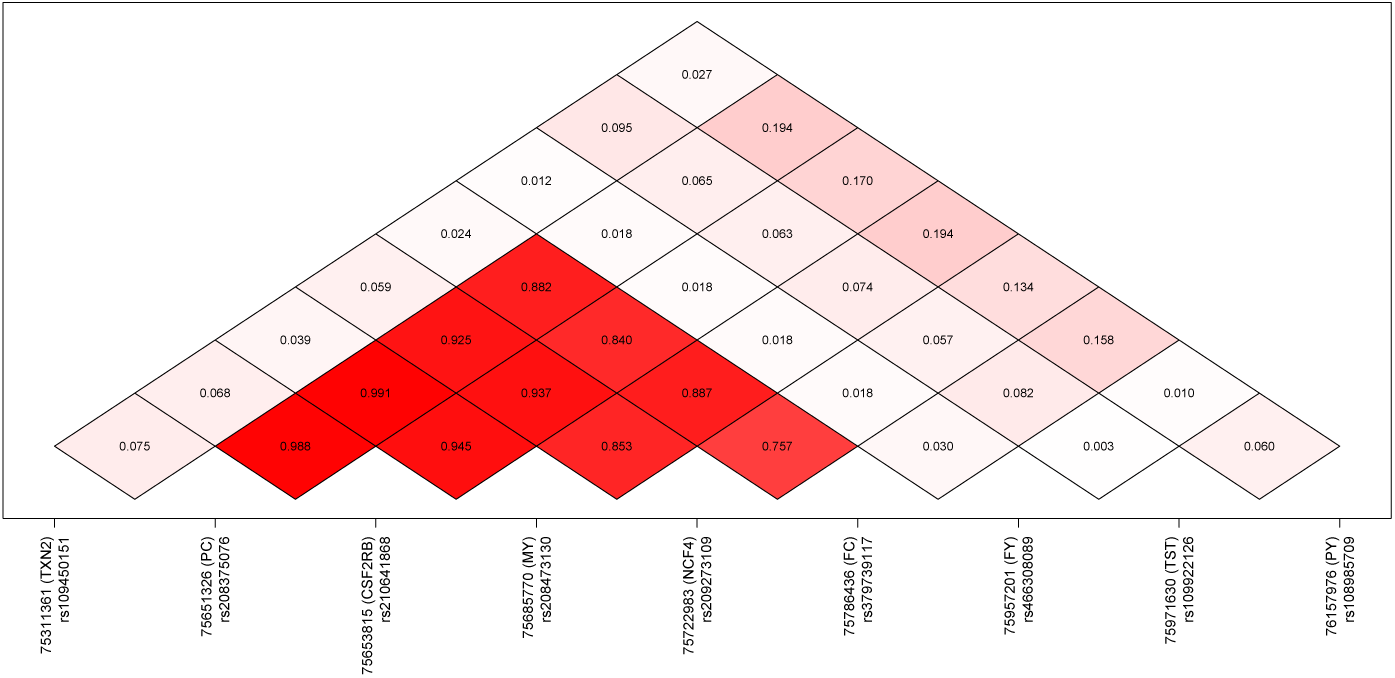
Linkage disequilibrium between the top tag variants for milk trait QTL and co-located gene expression QTL. Three genes with significant (P<5 × 10^−8^) eQTL are included, along with the *TST* [2] that have previously been proposed as a candidate causative gene at this locus.

### Evidence of multiple, differentially segregating QTL for milk yield and protein concentration

Examining panel A in Figure 1 (repeated in 5A) suggested that protein concentration might be influenced by two co-located but mechanistically independent QTL, as a number of markers that are not in strong LD with the top marker nevertheless exhibit very small p-values (*<* 1 × 10^−60^). To investigate this possibility, the top associated marker (rs208375076) was fitted as a fixed effect and the MLMA-LOCO analysis repeated using the residual, protein concentratiThe same pattern wason phenotype (Figure 5B). The new top marker (rs210293314) remained highly significant (P=1.30 × 10^−24^ after adjustment, 9.31 × 10^−41^ before adjustment), suggesting that it is tagging a different QTL. Adjusting the original protein concentration phenotype for rs210293314 and repeating the MLMA-LOCO analysis yielded the result shown in Figure 5C. Here, the most significant marker was rs208086849, a variant largely equivalent to the top rs208375076 marker from the original, unadjusted analysis (*R*^2^ = 0.999). These observations indeed suggest the presence of two QTL for milk protein percentage. This analysis was repeated with the MY phenotype (Figure 5D). This phenotype showed little evidence of a second co-locating QTL, where fitting the top associated marker (rs208473130) dropped the signal below the genome-wide significance threshold (P=1.36 × 10^−6^ for marker rs378861677; 5E). However, adjusting the MY phenotype by fitting rs378861677 and repeating the MLMA-LOCO analysis resulted in an increase in significance for the top marker rs208473130, from 6.64 × 10^−25^ to 8.63 × 10^−29^ (5F), suggesting there may indeed be an additional weak QTL, or the variant otherwise addresses some other confounding signal. The variants rs208086849 (from the PC analysis in the previous paragraph) and rs208473130 show very strong LD (*R*^2^ = 0.991), suggesting that both markers are in fact tagging the same QTL across the PC and MY traits. In contrast, variants rs210293314 (PC analysis above) and rs378861677 show moderate to weak LD (*R*^2^ = 0.332), suggesting the two signals tagged by these variants are genetically distinct.

**Figure 5.**
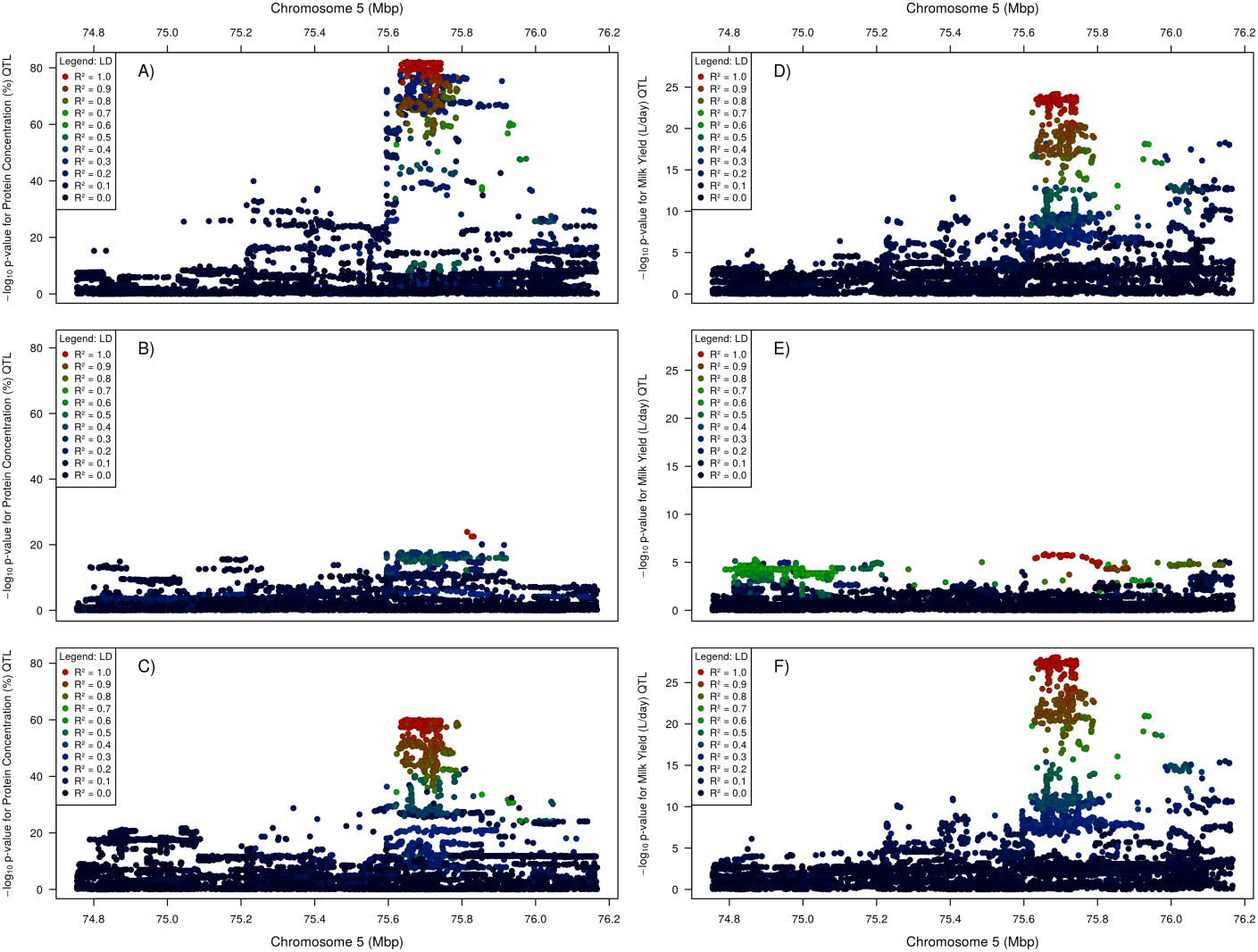
The effect of fitting the top variant on protein concentration (A–C) and milk yield (D–F) QTLs. The top panels (A & D) show the QTL with no marker adjustments fitted; the centre panels (B & E) show the QTL after fitting the top variant from the panel above; and the bottom panels (C & F) show the QTL after fitting the top variant from the centre panel above. The phenotypes were adjusted by fitting the following markers: B) rs208375076, C) rs210293314, D) rs208473130, E) rs378861677.

Since at least two differentially segregating QTL were apparent at the locus, it was possible that they were underpinned by different genes and/or molecular mechanisms. To assess whether the significant, co-locating *CSF2RB* and *NCF4* eQTL were themselves comprised of multiple, overlapping signals (i.e. multiple *cis*-eQTL driven by different regulatory elements), the top associated variants were fitted as fixed effects to the gene expression phenotypes, with analyses rerun as above. This yielded new top markers with p-values of 8.87 × 10^−5^ and 1.75 × 10^−4^ respectively, suggesting that the expression of these two genes, if influenced by multiple regulatory factors, were weak effects or impacts that were too heavily confounded by LD to differentiate clearly.

To look at how the eQTL might contribute to the multiple, co-locating PC QTL in comparative terms, the SNP-adjusted PC association results were used to calculate eQTL correlations, using the methodology described in the previous section. Notably, these analyses resulted in improved correlations with eQTL. The correlation between the *CSF2RB cis*-eQTL and the unadjusted PC phenotype was 0.754 (Figures 4 and 6A). However, using the phenotype adjusted for the marker rs210293314 yielded a correlation of 0.807 (6B). The same pattern was observed for the *NCF4* gene, where correlations improved from 0.691 to 0.843 (6C and D). Applying the same approach to milk yield phenotypes (unadjusted, and adjusted by marker rs37886167) gave similar results, albeit with marginal increases: correlations with the *CSF2RB* eQTL increased from 0.855 to 0.872, and correlations with the *NCF4* eQTL increased from 0.692 to 0.719.

**Figure 6.**
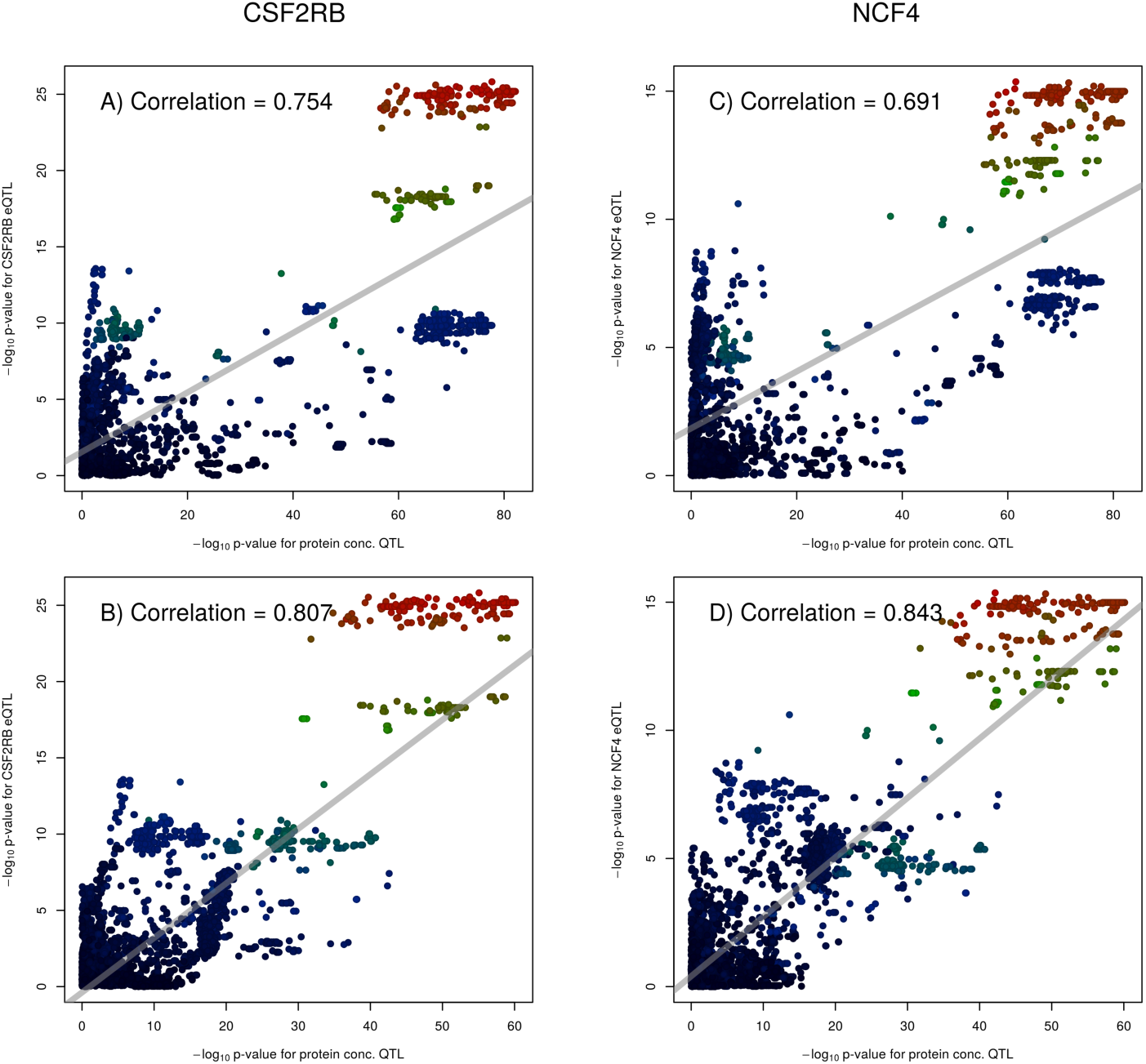
Correlations between eQTL and the co-located protein concentration QTL for the genes *CSF2RB* (left) and *NCF4* (right). Panels on the top row are plotted against the original protein concentration QTL (Figure 5A), while panels on the bottom row are plotted against the phenotype after fitting rs210293314 (Figure 5C).

To investigate the possibility that secondary, co-locating PC and/or MY QTL might be caused by protein-coding variants, all variants in strong LD (*R*^2^ *>* 0.9) with rs210293314 (secondary PC tag-SNP) or rs378861677 (secondary MY tag-SNP) were analysed using VEP as described previously. Of the 260 variants captured by this analysis, two missense SNPs were identified in conjunction with rs378861677, both mapping to exon two of the *MPST* gene: rs211170554 (p.Asp129Asn) with a SIFT score of 0.88 (predicted tolerated), and rs209917448 (p.Arg47Cys) with a SIFT score of 0.01 (predicted deleterious). In the absence of additional eQTL that might account for the secondary PC and MY signals, these results suggest a potential protein-coding-based mechanism for the MY effect at least.

### *CSF2RB* encodes a promiscuously RNA-edited transcript

Previous work [22] had identified four RNA editing sites mapping to the introns of the *CSF2RB* gene. Here, while manually examining RNAseq and WGS sequence reads mapping to the gene, a surprising number of additional RNA edits were apparent (see Materials and Methods). This included a total of 38 novel A-to-G variant sites present in the RNAseq data, yet absent from the whole genome sequence representing the nine cows for which both data sources were available. These sites were present in four clusters within the 3^*′*^ UTR (Figure 7), missed from our previously published genome-wide analysis [22] due to that analysis being based only on reference annotations that failed to capture the full length 3^*′*^UTR sequence evident using empirically derived gene structures from mammary RNAseq data. Because the ADAR genes responsible for adenosine-to-inosine editing (A-to-G in sequence reads) target double-stranded RNA [23, 24], we predicted the potential for the sequence surrounding the edited sites to form double-stranded RNA. The dot-plot shown in Figure 7 shows that, of the 38 edited sites (red dashed lines), 37 (97.4%) sit within regions of extended complementarity (diagonal black lines), thus having the potential to form double stranded secondary structures.

**Figure 7.**
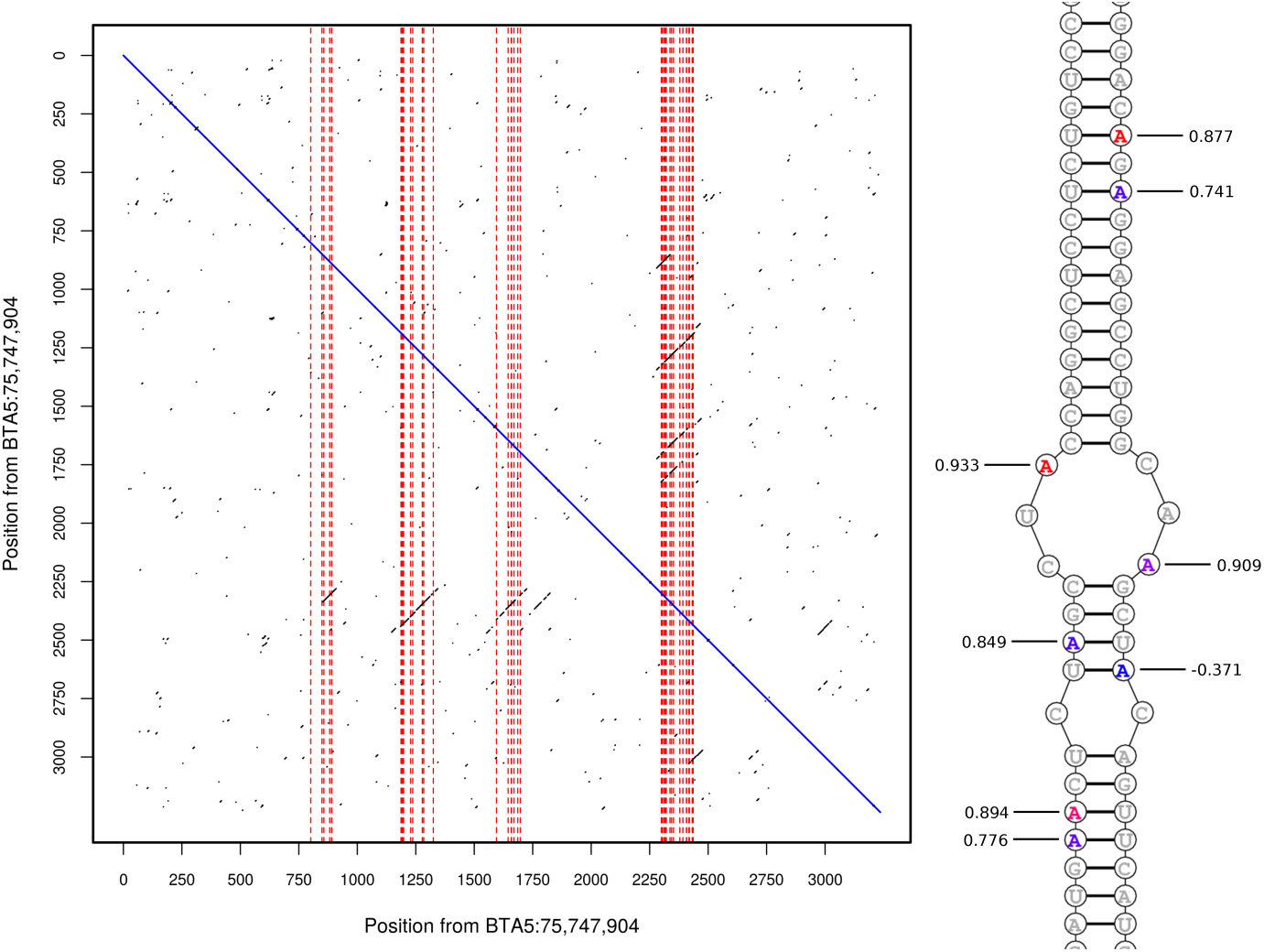
Left: dotplot of sequence from the *CSF2RB* 3^*′*^-UTR against its complement. Positions are given relative to BTA5:75,747,904. Black dots indicate that seven of the eleven surrounding nucleotides are complementary. Vertical dashed red lines indicate the locations of predicted RNA-editing sites. Sections of the region 2275–2452 are complementary to the regions 837–915, 1178–1350, 1591–1719, and 1757–1832, suggesting that the UTR is able to fold into multiple configurations. Right: the section of predicted double stranded sequence between 1184 and 1217 on the left strand (running upward), and 2411–2444 on the right strand (running downward). Edited sites are coloured based on the strength of the edQTL at that site, from blue (not significant) to red (max P=5.22 × 10^−26^). Sites are labelled with the correlation between the edQTL and the milk volume (MY) QTL after adjusting for marker rs208473130.

As recently reported, we have observed a proportion of RNA-edited bases to be genetically modulated for some sites [22]. To investigate potential genetic regulation of editing of *CSF2RB* transcripts, editing proportion phenotypes were generated (see Methods) for use in detecting RNA editing QTL (edQTL [22, 25]). Using the MLMA-LOCO method as applied for eQTL analysis described above, genome-wide significant edQTL (P<5 × 10^−8^) were identified for 18 of the 38 sites. Because RNA editing may impact gene expression by several different mechanisms [26, 27, 28], we investigated whether any edQTL were correlated with the eQTL for *CSF2RB*. One site, mapping to BTA5:75,750,220, had a correlation of 0.849 between the – log_10_ p-values of the edQTL and the eQTL. This edQTL was also strongly correlated with the *NCF4* eQTL (0.929).

As an extension to the hypothesis that edQTL might underlie changes in gene expression (i.e. eQTL), we reasoned that one or more of the milk phenotype QTL might also be impacted, as evidenced through correlation. Investigating this hypothesis, we observed correlations *>* 0.707 between edQTL and the fat concentration, protein concentration, and protein yield phenotypes (Table 5). In addition, we found very strong correlations (*>* 0.9) between two edQTL (sites BTA5:75,749,101 and BTA5:75,750,335) and the milk yield phenotype after adjusting for the genotype of marker rs208473130 (yield QTL illustrated in Figure 5E, correlations in Figure 7). A strong correlation (0.822) was also detected between the edQTL for site BTA5:75,748,760 and the protein concentration QTL after adjusting for marker rs208375076 (PC QTL illustrated in Figure 5B). Like analyses of candidate proteincoding variants, these results suggest other alternative (and likely overlapping) mechanisms that may account for the multiple QTL segregating at the chromosome 5 locus.

**Table 1.**
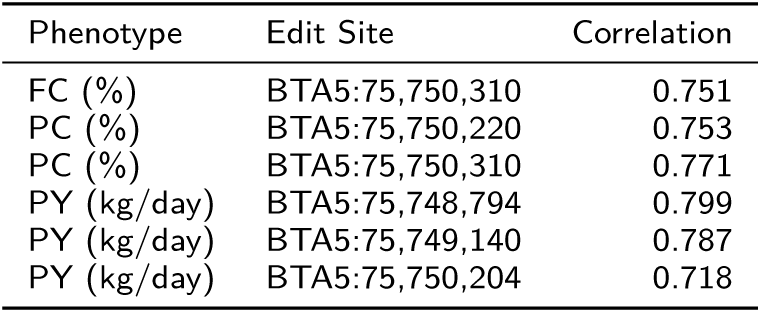
Pearson correlations between the – log_10_ p-values for milk trait QTL and edQTL for sites mapping to the 3^*′*^-UTR of *CSF2RB*. Only sites and phenotypes where the correlation exceeded 0.707 (*R*^2^ *>* 0.5) are shown.

### Hypervariability at the *CSF2RB* locus presents an abundance of candidate causative variants

Manual examination of the WGS alignments at the locus also revealed read depth anomalies at approximately BTA5:75,781,300–75,782,800. This analysis revealed a suspected 1.5 kbp deletion variant, located between the *CSF2RB* and *TEX33* genes (downstream of the 3^*′*^ UTRs of both genes given a ‘tail to tail’ orientation). To attempt to derive genotypes for this variant, the copy number at this site was estimated for 560 whole genome sequenced cattle using CNVnator 0.3 [29]. The resulting estimates of copy number formed a trimodal distribution (Figure 8A), suggestive of a biallelic variant that could be assumed to be inherited in a Mendelian fashion [30]. Although one pseudogene maps to the region (LOC788541 60S ribosomal protein L7), the deleted segment appeared otherwise devoid of noteworthy genomic features.

**Figure 8.**
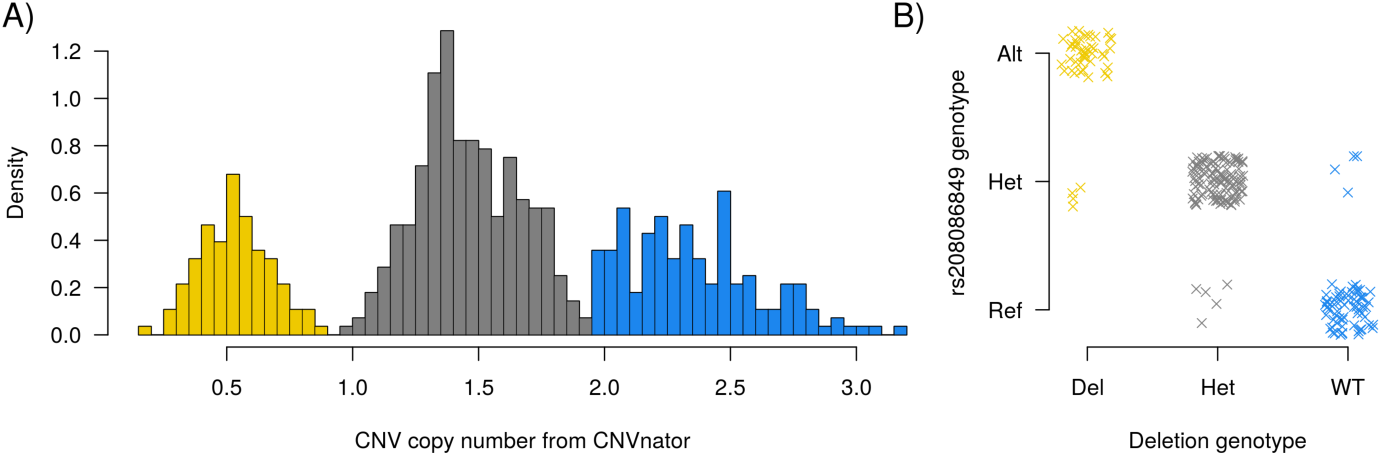
A) Histogram of copy number genotype calls of 560 animals from CNVnator. Copy numbers follow a trimodal distribution, suggesting that the variant is bialleleic. Genotype classes are coloured gold (homozygous deletion), grey (heterozygous) and blue (homozygous wild-type). B) deletion variant genotypes plotted against the genotypes of the rs208086849 variant. The two variants are in strong LD (R^2^=0.887). Points are jittered to increase visibility.

To investigate the candidacy of the deletion as a potential causative variant for one or more of the QTL in the region, genotypes were called from CNVnator copy number predictions (Materials and Methods), and the LD (R^2^) between the deletion and top QTL variants investigated. Strong LD (0.887) was observed with the top markers for the MY (rs208473130) and PC (rs208375076) phenotypes, as well as with rs208086849, the top variant for the PC phenotype after adjusting for the secondary QTL (Figure 8B). A slightly lower LD score was observed for the FC phenotype (R^2^=0.807). The deletion allele was more frequent than the reference allele in the NZ dairy population (deletion=0.547).

The strong LD of the *∼* 1.5 kbp deletion with key QTL tag variants qualified the variant as a potential candidate for these QTL, so we then imputed the variant into the association analysis population to test for association directly. Using the same MLMA-LOCO analysis method applied for other variants, significant associations (P<5 × 10^−8^) were observed for the PC (P=7.30 × 10^−71^), FC (P=1.08 × 10^−30^), and MY (1.18 × 10^−18^) phenotypes. Although highly significant, when ranking all variants by p-value, the deletion variant never placed higher than the 400^th^ most significant; however, given the very large number of associated variants at the region generally (>800 in the top 20 orders of magnitude for PC), and the fact that some of the read-depth-based genotype calls may be erroneous, the deletion remains a plausible candidate variant for future consideration of these QTL.

## Discussion

### Milk phenotype QTL

We report QTL mapping of a chromosome 5 locus for several milk yield and composition phenotypes, with a diversity of gene expression and RNA editing QTL that could underpin these effects. We note in particular that some phenotypes exhibit multiple QTL, likely with distinct genetic causes. The fat and protein concentration QTL are both in high LD with the milk yield QTL, suggesting that these effects may be mediated by changes in the total volume of milk produced without concomitant changes in fat or protein production. The fat and protein yield QTL are not in LD with either each other or with milk yield. However, these two QTL are less significant than the others by many orders of magnitude (see Table 1), suggesting that the lack of LD may be due to insufficient power in the data set to identify reproducible tag variants.

### Candidate causative genes

Several candidate causative genes have been previously proposed to underlie lactation effects at this locus, and based on the work presented here, we propose that one or both of the *CSF2RB* and *NCF4* genes are the likely candidates, with a predicted deleterious variant in the *MPST* gene also providing a potential candidate for a secondary effect milk yield QTL.

The *CSF2RB* gene (ENSBTAG00000009064) encodes the common beta chain of the receptors for GM-CSF, interleukin-3, and interleukin-5, cytokines that are involved in regulating the proliferation and differentiation of hematopoietic cells [31]. Granulocyte-macrophage colony-stimulating factor (GM-CSF) is produced in the mammary gland by alveolar macrophages [32] where it enhances the bactericidal activity of milk neutrophils [33]. These receptors form a link in the JAK-STAT signalling pathway, operating via JAK2 and STAT5 [34]. The STAT5 proteins, especially STAT5A, are important for enabling mammopoiesis and lactogenesis [35, 36] and directly bind the gamma-interferon-activating sequence (GAS) found in the promoters of milk proteins such as beta-casein, [37], beta-lactoglobulin [38], and whey acidic protein in mice [38]. The importance of this pathway is further evidenced by associations for milk production traits observed at the *STAT5* locus [39, 40, 41]. Although the relevant ligands and subunits with which *CSF2RB* forms complexes are unknown in the current context, mutations that impact downstream interactions with STAT5 proteins could be assumed to impact milk production/composition phenotypes.

The *NCF4* gene (ENSBTAG00000007531) encodes neutrophil cytosolic factor 4, which forms the p40-phox subunit of the NADPH oxidase enzyme complex [42]. This enzyme produces superoxide 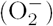, a reactive oxygen species produced in phagocytic cells during the respiratory burst [43], intended to kill invading fungi and bacteria [44]. *NCF4* has been shown to be upregulated in mastitic mammary glands [45], and two SNPs mapping to the *NCF4* gene have been associated with elevated somatic cell scores (SCS) [45, 46], a trait that is used as a surrogate phenotype for mastitis in dairy animals. As cows suffering from mastitis produce smaller volumes of milk than healthy cows [47], this provides a possible mechanism by which *NCF4* could influence milk production. A more appealing mechanism is one that involves *CSF2RB* or *NCF4* outside of a pathogen response context however, given that the locus is better known for its impacts on milk production and composition in the absence of overt mammary infection.

Both the *CSF2RB* and *NCF4* eQTL were correlated with the MY QTL, with the former giving stronger correlations (*r* = 0.849 compared to 0.682). Lower correlations were observed between the two eQTL and the PC QTL (*r* = 0.754 and 0.691), however, removing one of the two apparent signals at this locus by fitting the rs210293314 marker to the PC phenotype increased correlations for both candidate genes. As no other genes showed similar patterns of co-association, we consider one or more of these genes to be the best candidates at this locus. The *CSF2RB* gene was expressed very strongly in mammary samples (TPM=80.1), and by comparison, at a much higher level than *NCF4* (TPM=0.51). This observation suggests a critical role for *CSF2RB*-mediated signalling in lactation, and given the plausible biological linkages of *CSF2RB* to these processes (via JAK-STAT signalling), we favour *CSF2RB* as the more likely of these two candidates.

The *TST* gene (ENSBTAG00000030650) was recently proposed by Pausch et al. [2] as a candidate for milk fat and protein percentage QTL at *∼* 75–76 Mbp on chromosome 5. *TST* encodes thiosulfate sulfurtransferase, also known as rhodanese, a mitochondrial enzyme that catalyses the conversion of cyanide plus thiosulfate into thiocyanate plus sulfite [48]. It has also been shown that the rhodanese enzyme (in misfolded form) can bind with 5S-rRNA, enabling its import into the mitochondria [49]. There appears to be limited literature implicating *TST* in mammary development and milk production, and given that the gene maps downstream of peak association in our dataset, and has no prominent eQTL by which to mediate these effects, a role for this gene seems unlikely for QTL in the NZ population. This does not preclude the involvement of the gene in other populations, however, we submit that the most parsimonious hypothesis is that these QTL are shared across populations, at least partially underpinned by regulatory variants modulating the expression of the *CSF2RB* gene.

### RNA editing and edQTL

We previously [10] reported four RNA editing sites mapping to the *CSF2RB* gene, one of which (BTA5:75,739,106) showed a significant edQTL (smallest P=6.68 × 10^−13^). This site exhibited only modest correlations with the *CSF2RB* eQTL, or with the milk yield or composition QTL [10].

In the current paper, we report the discovery of an additional 38 RNA-editing sites mapping to the 3^*′*^-UTR of *CSF2RB*. These sites were not identified in the previous work as they map approximately 3 kbp downstream of the gene structure based on the Ensembl reference annotation. Two of the novel sites, BTA5:75,749,101 and BTA5:75,750,335, exhibited edQTL with correlations exceeding 0.9 with the milk yield QTL after adjusting for marker rs208473130. The correlation between the *CSF2RB* eQTL and the same milk QTL was -0.173, suggesting that, if the lactation effects indeed derive from an RNA-editing-based mechanism, that this mechanism is not wholly reflected by the gene expression data used to quantify the eQTL effects.

## Conclusions

We have examined a previously implicated chromosome 5 locus for milk yield and composition traits, and identified highly significant QTL for milk yield, protein concentration, and fat concentration. Using a large mammary RNA sequence resource, we have conducted eQTL mapping of the locus and show that expression of *CSF2RB*, a highly expressed gene that signals through pathways important to mammary development and lactation, appears to be responsible for these effects. RNA editing sites were also discovered in the 3^*′*^-UTR of *CSF2RB*, and edQTL for two of these are correlated with one of two co-located yet differentially segregating milk yield QTL, which was also in strong LD with a predicted deleterious missense variant in the *MPST* gene. These results highlight the pleiotropic nature of the *CSF2RB* gene, and showcase the mechanistic complexity of a locus that will require further statistical and functional dissection to catalogue the full multiplicity of effects.

## Methods

### Genotyping and phenotyping

All cows that had been genotyped using the GGP LDv3 or LDv4 chips for which herd test phenotypes were also available were targeted for the current study (N=29,350). Animals representing these platforms were targeted since, based on preliminary sequence-based association analyses not reported here, these panels had been enriched with 365 polymorphisms identified as tag-variants of the chromosome 5 lactation QTL (spanning a region from 74.8–76.2 Mbp; Additional File 1). These variants included 30 SNPs from the Illumina BovineSNP50 chip (50k), added to assist with imputation of the region and create equivalence with that platform for other analysis applications. Tag-variants were targeted as custom content using a scheme that attempted to genotype sites in both orientations (two primers per site), resulting in 341 custom markers on the LDv3 chip, and 342 on the LDv4 chip for this locus. Phenotypes were calculated from their herd-test records for the three yield traits plus fat and protein concentration in milk. These phenotypes were generated using herd-test data from the first lactation, adjusted using an ASReml-R [50] model with birth year, age at calving, breed, and heterosis as linear covariates, stage of lactation as a fixed effect, season/herd as an absorbed fixed effect, and animal as a random effect. Herd test records were sampled using Fourier-transform infrared spectroscopy on a combination of Milkoscan FT6000 (FOSS, Hillerød, Denmark) and Bentley FTS (Bentley, Chaska, USA) instruments.

### Imputation and association analyses

Genotypes for 29,350 animals were imputed to WGS resolution in the window of interest using Beagle 4 [15] as described previously [51, 9]. Briefly, a reference population of 565 animals, comprising Holstein-Friesians, Jerseys, and cross-bred cattle, was sequenced using the Illumina HiSeq 2000 instrument to yield 100 bp reads. Read mapping to the UMD 3.1 bovine reference genome was conducted using BWA MEM 0.7.8 [52], followed by variant calling using GATK HaplotypeCaller 3.2 [53]. Variants were phased using Beagle 4 [15], and those with poor phasing metrics (allelic R^2^ < 0.95) were excluded, yielding 12,867 variants. Quality control filtering to further remove variants with MAF<0.01% (N=673) or Hardy-Weinberg Equilibrium p-values <1 × 10^−30^ (N=461) resulted in a final set of 11,733 variants. As described above, the imputation window was enriched for custom, physically genotyped variants on the GGP-LDv3/4 chips, markedly increasing the scaffold density at this location.

Imputed genotypes for 639,822 autosomal markers on the Illumina BovineHD SNP-chip were used to calculate a genomic relationship matrix (GRM) for the 29,350 animals of interest, using GCTA [54, 16]. The imputation step also used Beagle 4 software, leveraging a BovineHD-genotyped reference population of 3,389 animals. Heritabilities for all phenotypes were calculated using this GRM with the REML option in GCTA. A leave-one-chromosome-out (LOCO) GRM was also created excluding chromosome 5, and used in combination with the imputed variant set and phenotypes to perform a mixed linear model analysis (MLMA-LOCO) [55] using GCTA.

### RNAseq, gene expressions and eQTL

RNAseq data from lactating mammary gland biopsies representing 357 cows was generated as described previously [41]. Briefly, samples were sequenced using Illumina HiSeq 2000 instruments, yielding 100 bp paired-end reads. These were mapped to the UMD 3.1 reference genome using TopHat2 [56]. Stringtie software [18] was used to quantify gene expression values for genes mapping to the window BTA5:75–76Mbp, yielding fragments per kilobase of transcript per million mapped reads (FPKM) and transcripts per million (TPM) [57] metrics. These calculations used gene models defined by the Ensembl gene build (release 81). Gene expression levels were also processed using the variance-stabilising transformation (VST) function implemented in the Bioconductor package DESeq [21] to produce expression data suitable for analysis using linear models.

WGS-resolution genotypes were imputed using the same WGS sequence reference described above in conjunction with a mixture of genotype panels (see Methods in [41]) for the 357 cows, yielding 12,825 variants in the window BTA5:74.6–76.2Mbp. Filtering to remove variants with >5% missing genotypes (N=36) or MAF <0.5% (N=1,643) resulted in a final set containing 11,146 variants. VST-transformed gene expressions were analysed for genes with FPKM > 0.1, using the GCTA MLMA-LOCO method described above. The GRM was calculated using physically genotyped variants from the BovineHD SNP chip for 337 cows, and imputed BovineHD genotypes for the remaining twenty cows based on an Illumina SNP50 platform scaffold.

### RNA-editing site discovery and edQTL

RNA editing in the 3^*′*^-UTR of the *CSF2RB* gene was investigated in the nine discovery set animals from [22], where these animals had been previously sequenced using both RNAseq and WGS methodologies. Editing sites were identified using custom scripts [22] and by manual inspection of WGS and RNAseq BAM files for each animal. Sites were considered to represent RNA edits where: i) an A-to-G variant was present in the RNAseq reads, but was absent from the WGS reads, and ii) had at least five reads containing ‘G’ at the position in every animal. This yielded 38 candidate edited sites. Following the recommendations of Ramaswami et al 2012 for non-*Alu* sites, the 38 candidate sites were examined for the presence of 5^*′*^ mismatches, simple repeats, homopolymer runs *≥* 5 bp, or splice junctions within 4 bp; however, none of the candidates were impacted by these filters, and all 38 were retained for further analyses.

Having determined the positions of variant sites, the rate of editing at each site was quantified in the larger ‘quantification set’ of 353 cows [22] with RNA editing phenotypes for each site generated by transforming editing proportions using the logit function. RNA editing QTL discovery was performed using these phenotypes by performing MLMA-LOCO, incorporating the same GRM and imputed WGS genotypes used for eQTL discovery (N=353 animals).

RNA secondary structure around the edited sites was predicted using dot-plots as described by [22]. Sequence containing all 38 edited sites was extracted, along with 800 bp upstream and downstream. The sequence was then plotted against its complement, with dots placed where at least 11 of 15 nucleotides surrounding a point were complementary. Diagonal lines in the resulting plot indicate regions of extended complementarity, which therefore have the potential to form double-stranded secondary structures.

### Copy number variant genotyping and imputation

Manual examination of the WGS BAM files suggested the presence of a copy-number variant (CNV) located downstream of *CSF2RB*, mapping to BTA5:75,781,300– 75,782,800. Copy numbers were estimated from WGS reads for each of 560 cattle using the software package CNVnator version 0.3 [29], based on sequence read depth. Thresholds for genotype calling of the CNV were decided based on the histogram of the trimodal distribution of the copy number (CN) estimates, where homozygous deletion was called where CN<0.95, heterozygous 0.95 <= CN < 1.95, and homozygous wild type when CN >= 1.95. CNV genotypes were imputed into a larger population (N=29,350), for use in association analyses, using Beagle version 4.1 [59], using the reference population of 560 cattle described above. Combining the reference genotype calls with the imputed population yielded a set of 31,950 animals for use in MLMA-LOCO analyses, as described above.

## List of abbreviations

CNV: Copy-number variant
FC: Milk fat concentration
FY: Milk fat yield
GRM: Genomic relationship matrix
LOCO: Leave one chromosome out
MY: Milk yield
PC: Milk protein concentration
PY: Milk protein yield
QTL: Quantitative trait locus
RNAseq: RNA sequence data
WGS: Whole genome sequence
YD: yield deviation

## Declarations

### Ethics approval

All animal experiments were conducted in strict accordance with the rules and guidelines outlined in the New Zealand Animal Welfare Act 1999. Most data were generated as part of routine commercial activities outside the scope of that requiring formal committee assessment and ethical approval (as defined by the above guidelines). These animals were located in commercial dairy herds around New Zealand, with approval given to tissue sample for genetic analyses. For the mammary tissue RNA sequencing biopsy experiment, samples were obtained in accordance with protocols approved by the Ruakura Animal Ethics Committee, Hamilton, New Zealand (approval AEC 12845). These cows were situated on a research farm and permission was sought and obtained to biopsy mammary tissue from the owner of these animals (AgResearch, NZ). No animals were sacrificed for this study.

### Consent for publication

Not applicable.

### Availability of data and material

Sequence data representing the chromosome 5 locus of interest are available for download and have been added to the sequence read archive (SRA=SRP159443). Whole genome and mammary transcriptome sequences representing the nine animals used for RNA-edit discovery have also been uploaded to SRA (SRP136662). The GRM, phenotype data, and imputed sequence-based genotype data representing the animals used for lactation trait association analyses (N=29,350), and RNAseq eQTL/edQTL analyses (N=357), are also available for download and can be accessed through the Dryad database portal (Dryad DOI=TBA).

### Authors’ contributions

TJL performed most of the bioinformatic and statistical analyses with help from KT and CC; TJL, SRD, RGS, and MDL conceived of the study and experiments; SRD, RGS, RJS, and MDL were involved in supervision of the project; TJL and MDL wrote the manuscript. All authors have read and approved the manuscript.

## Acknowledgements

The authors would like to acknowledge S. Morgan and staff at DairyNZ Ltd. (Hamilton, New Zealand), and Phil McKinnon, Ali Cullum and staff at AgResearch (Hamilton, New Zealand) for facilitating mammary tissue sampling of lactating animals. We also wish to acknowledge New Zealand Genomics Limited (NZGL) and the University of Auckland Centre for Genomics, Proteomics, and Metabolomics for RNA preparation and sequencing, as well as both the Australian Genome Research Facility (AGRF) and Illumina FastTrack for both RNA and genomic DNA sequencing.

## Additional Files

### Additional File 1.xlsx

Polymorphisms targeted as custom content for the GGP LDv3 and LDv4 chips. These were identified as tag variants of chromosome 5 lactation QTL in the window 74.8–76.2 Mbp.

## Author details

^1^ Research & Development, Livestock Improvement Corporation, Ruakura Road, Hamilton, NZ. ^2^ School of Biological Sciences, University of Auckland, Symonds Street, Auckland, NZ.

## References

1. Raven LA, Cocks BG, Kemper KE, Chamberlain AJ, Vander Jagt CJ, Goddard ME, et al. Targeted imputation of sequence variants and gene expression profiling identifies twelve candidate genes associated with lactation volume, composition and calving interval in dairy cattle. Mammalian genome. 2016;27:81–97. doi:10.1007/s00335-015-9613-8.

2. Pausch H, Emmerling R, Gredler-Grandl B, Fries R, Daetwyler HD, Goddard ME. Meta-analysis of sequence-based association studies across three cattle breeds reveals 25 QTL for fat and protein percentages in milk at nucleotide resolution. BMC Genomics. 2017;18:853. doi:10.1186/s12864-017-4263-8.

3. Wang T, Chen Y4P, MacLeod IM, Pryce JE, Goddard ME, Hayes BJ. Application of a Bayesian non-linear model hybrid scheme to sequence data for genomic prediction and QTL mapping. BMC Genomics. 2017;18:618. doi:10.1186/s12864-017-4030-x.

4. Calus M, Goddard M, Wientjes Y, Bowman P, Hayes B. Multibreed genomic prediction using multitrait genomic residual maximum likelihood and multitask Bayesian variable selection. Journal of dairy science. 2018;101. doi:10.3168/jds.2017-13366.

5. Grisart B, Coppieters W, Farnir F, Karim L, Ford C, Berzi P, et al. Positional candidate cloning of a QTL in dairy cattle: identification of a missense mutation in the bovine DGAT1 gene with major effect on milk yield and composition. Genome research. 2002;12(2):222–231. doi:10.1101/gr.224202.

6. Cohen-Zinder M, Seroussi E, Larkin DM, Loor JJ, Everts-van der Wind A, Lee JH, et al. Identification of a missense mutation in the bovine ABCG2 gene with a major effect on the QTL on chromosome 6 affecting milk yield and composition in Holstein cattle. Genome Research. 2005;15(7):936–944. doi:10.1101/gr.3806705.

7. Blott S, Kim JJ, Moisio S, Schmidt-Küntzel A, Cornet A, Berzi P, et al. Molecular dissection of a quantitative trait locus: a phenylalanine-to-tyrosine substitution in the transmembrane domain of the bovine growth hormone receptor is associated with a major effect on milk yield and composition. Genetics. 2003;163(1):253–266.

8. Kemper K, Littlejohn M, Lopdell T, Hayes B, Bennett L, Williams R, et al. Leveraging genetically simple traits to identify small-effect variants for complex phenotypes. BMC genomics. 2016;17:858. doi:10.1186/s12864-016-3175-3.

9. Littlejohn MD, Tiplady K, Fink TA, Lehnert K, Lopdell T, Johnson T, et al. Sequence-based association analysis reveals an MGST1 eQTL with pleiotropic effects on bovine milk composition. Scientific Reports. 2016;6:25376. doi:10.1038/srep25376.

10. Lopdell T, Tiplady K, Littlejohn M. Using RNAseq data to improve genomic selection in dairy cattle. In: Proceedings of the World Congress on Genetics Applied to Livestock Production. vol. 11. Auckland, New Zealand; 2018. p. 49.

11. Meuwissen T, Hayes B, Goddard M. Prediction of total genetic value using genome-wide dense marker maps. Genetics. 2001;157(4):1819–1829.

12. Habier D, Fernando RL, Kizilkaya K, Garrick DJ. Extension of the Bayesian alphabet for genomic selection. BMC bioinformatics. 2011;12(1):186. doi:10.1186/1471-2105-12-186.

13. Kemper KE, Reich CM, Bowman PJ, Vander Jagt CJ, Chamberlain AJ, Mason BA, et al. Improved precision of QTL mapping using a nonlinear Bayesian method in a multi-breed population leads to greater accuracy of across-breed genomic predictions. Genetics Selection Evolution. 2015;47:29. doi:10.1186/s12711-014-0074-4.

14. Kemper K, Hayes B, Daetwyler H, Goddard M. How old are quantitative trait loci and how widely do they segregate? Journal of Animal Breeding and Genetics. 2015;132(2):121–134. doi:10.1111/jbg.12152.

15. Browning BL, Browning SR. A unified approach to genotype imputation and haplotype-phase inference for large data sets of trios and unrelated individuals. The American Journal of Human Genetics. 2009;84(2):210–223. doi:10.1016/j.ajhg.2009.01.005.

16. Yang J, Lee SH, Goddard ME, Visscher PM. GCTA: a tool for genome-wide complex trait analysis. The American Journal of Human Genetics. 2011;88(1):76–82. doi:10.1016/j.ajhg.2010.11.011.

17. McLaren W, Gil L, Hunt SE, Riat HS, Ritchie GR, Thormann A, et al. The Ensembl Variant Effect Predictor. Genome biology. 2016;17:122. doi:10.1186/s13059-016-0974-4.

18. Pertea M, Pertea GM, Antonescu CM, Chang TC, Mendell JT, Salzberg SL. StringTie enables improved reconstruction of a transcriptome from RNA-seq reads. Nature biotechnology. 2015;33(3):290. doi:10.1038/nbt.3122.

19. Yue F, Cheng Y, Breschi A, Vierstra J, Wu W, Ryba T, et al. A comparative encyclopedia of DNA elements in the mouse genome. Nature. 2014;515(7527):355. doi:10.1038/nature13992.

20. Trinklein ND, Aldred SF, Hartman SJ, Schroeder DI, Otillar RP, Myers RM. An abundance of bidirectional promoters in the human genome. Genome research. 2004;14(1):62–66. doi:10.1101/gr.1982804.

21. Anders S, Huber W. Differential expression analysis for sequence count data. Genome biology. 2010;11(10):R106. doi:10.1186/gb-2010-11-10-r106.

22. Lopdell TJ, Couldrey C, Tiplady K, Davis SR, Snell RG, Harris BL, et al. Widespread *cis*-regulation of RNA-editing in a large mammal. bioRxiv. 2018;p. 304220. doi:10.1101/304220.

23. Lehmann KA, Bass BL. The importance of internal loops within RNA substrates of ADAR1. Journal of molecular biology. 1999;291(1):1–13. doi:10.1006/jmbi.1999.2914.

24. Lehmann KA, Bass BL. Double-stranded RNA adenosine deaminases ADAR1 and ADAR2 have overlapping specificities. Biochemistry. 2000;39(42):12875–12884. doi:10.1021/bi001383g.

25. Ramaswami G, Deng P, Zhang R, Carbone MA, Mackay TF, Li JB. Genetic mapping uncovers *cis*-regulatory landscape of RNA editing. Nature communications. 2015;6:8194. doi:10.1038/ncomms9194.

26. Wang Q, Hui H, Guo Z, Zhang W, Hu Y, He T, et al. ADAR1 regulates ARHGAP26 gene expression through RNA editing by disrupting miR-30b-3p and miR-573 binding. RNA. 2013;19(11):1525–1536. doi:10.1261/rna.041533.113.

27. Brümmer A, Yang Y, Chan TW, Xiao X. Structure-mediated modulation of mRNA abundance by A-to-I editing. Nature communications. 2017;8(1):1255. doi:10.1038/s41467-017-01459-7.

28. Prasanth KV, Prasanth SG, Xuan Z, Hearn S, Freier SM, Bennett CF, et al. Regulating gene expression through RNA nuclear retention. Cell. 2005;123(2):249–263. doi:DOI:10.1016/j.cell.2005.08.033.

29. Abyzov A, Urban AE, Snyder M, Gerstein M. CNVnator: an approach to discover, genotype, and characterize typical and atypical CNVs from family and population genome sequencing. Genome research. 2011;21(6):974–984. doi:10.1101/gr.114876.110.

30. Couldrey C, Keehan M, Johnson T, Tiplady K, Winkelman A, Littlejohn M, et al. Detection and assessment of copy number variation using PacBio long-read and Illumina sequencing in New Zealand dairy cattle. Journal of dairy science. 2017;100(7):5472–5478. doi:10.3168/jds.2016-12199.

31. Miyajima A, Mui A, Ogorochi T, Sakamaki K. Receptors for granulocyte-macrophage colony-stimulating factor, interleukin-3, and interleukin-5. Blood. 1993;82(7):1960–1974.

32. Ito T, Kodama M. Demonstration by reverse transcription-polymerase chain reaction of multiple cytokine mRNA expression in bovine alveolar macrophages and peripheral blood mononuclear cells. Research in veterinary science. 1996;60(1):94–96. doi:10.1016/S0034-5288(96)90140-X.

33. Alluwaimi AM. The cytokines of bovine mammary gland: prospects for diagnosis and therapy. Research in veterinary science. 2004;77(3):211–222. doi:10.1016/j.rvsc.2004.04.006.

34. Mak TW, Saunders ME. Cytokines and Cytokine Receptors. In: The Immune Response: Basic and Clinical Principles. Academic Press; 2006. p. 463–516. doi:10.1016/B978-012088451-3.50019-3.

35. Liu X, Robinson GW, Wagner KU, Garrett L, Wynshaw-Boris A, Hennighausen L. Stat5a is mandatory for adult mammary gland development and lactogenesis. Genes & development. 1997;11(2):179–186. doi:10.1101/gad.11.2.179.

36. Gallego MI, Binart N, Robinson GW, Okagaki R, Coschigano KT, Perry J, et al. Prolactin, growth hormone, and epidermal growth factor activate Stat5 in different compartments of mammary tissue and exert different and overlapping developmental effects. Developmental biology. 2001;229(1):163–175. doi:10.1006/dbio.2000.9961.

37. Schmitt-Ney M, Doppler W, Ball R, Groner B. Beta-casein gene promoter activity is regulated by the hormone-mediated relief of transcriptional repression and a mammary-gland-specific nuclear factor. Molecular and Cellular Biology. 1991;11(7):3745–3755. doi:10.1128/MCB.11.7.3745.

38. Liu X, Robinson GW, Gouilleux F, Groner B, Hennighausen L. Cloning and expression of Stat5 and an additional homologue (Stat5b) involved in prolactin signal transduction in mouse mammary tissue. Proceedings of the National Academy of Sciences. 1995;92(19):8831–8835. doi:10.1073/pnas.92.19.8831.

39. Selvaggi M, Albarella S, Dario C, Peretti V, Ciotola F. Association of *STAT5A* Gene Variants with Milk Production Traits in Agerolese Cattle. Biochemical genetics. 2017;55(2):158–167. doi:10.1007/s10528-016-9781-6.

40. Ratcliffe L, Mullen M, McClure M, McClure J, Kearney F. 190 Single nucleotide polymorphisms in the signal transducer and regulator of transcription (STAT) genes are associated with milk production, milk composition, and fertility traits in Holstein Friesian cattle. Journal of Animal Science. 2017;951(suppl_4):94–94. doi:10.2527/asasann.2017.190.

41. Lopdell TJ, Tiplady K, Struchalin M, Johnson TJ, Keehan M, Sherlock R, et al. DNA and RNA-sequence based GWAS highlights membrane-transport genes as key modulators of milk lactose content. BMC Genomics. 2017;18(1):968. doi:10.1186/s12864-017-4320-3.

42. Leusen JH, Verhoeven AJ, Roos D. Interactions between the components of the human NADPH oxidase: a review about the intrigues in the phox family. Frontiers in Bioscience. 1996;1:d72–d90.

43. Decoursey T, Ligeti E. Regulation and termination of NADPH oxidase activity. CMLS Cellular and Molecular Life Sciences. 2005;62(19-20):2173–2193. doi:10.1007/s00018-005-5177-1.

44. Heyworth PG, Cross AR, Curnutte JT. Chronic granulomatous disease. Current opinion in immunology. 2003;15(5):578–584. doi:10.1016/S0952-7915(03)00109-2.

45. Ju Z, Wang C, Wang X, Yang C, Sun Y, Jiang Q, et al. Role of an SNP in alternative splicing of bovine NCF4 and mastitis susceptibility. PLoS one. 2015;10(11):e0143705. doi:10.1371/journal.pone.0143705.

46. Ju Z, Wang C, Wang X, Yang C, Zhang Y, Sun Y, et al. The effect of the SNP g.18475 A>G in the 3^<sup>1 </sup>^UTR of NCF4 on mastitis susceptibility in dairy cattle. Cell Stress and Chaperones. 2018;23:385–391.

47. Lescourret F, Coulon J. Modeling the impact of mastitis on milk production by dairy cows. Journal of Dairy Science. 1994;77(8):2289–2301. doi:10.3168/jds.S0022-0302(94)77172-1.

48. Cipollone R, Ascenzi P, Tomao P, Imperi F, Visca P. Enzymatic detoxification of cyanide: clues from *Pseudomonas aeruginosa* Rhodanese. Journal of molecular microbiology and biotechnology. 2008;15(2-3):199–211. doi:10.1159/000121331.

49. Smirnov A, Comte C, Mager-Heckel AM, Addis V, Krasheninnikov IA, Martin RP, et al. Mitochondrial enzyme rhodanese is essential for 5 S ribosomal RNA import into human mitochondria. Journal of Biological Chemistry. 2010;285(40):30792–30803. doi:10.1074/jbc.M110.151183.

50. Butler D, Cullis B, Gilmour A, Gogel B. ASReml-R reference manual: mixed models for S language. Brisbane, Queensland, Australia: Queensland Government; 2009.

51. Littlejohn MD, Henty KM, Tiplady K, Johnson T, Harland C, Lopdell T, et al. Functionally reciprocal mutations of the prolactin signalling pathway define hairy and slick cattle. Nature communications. 2014;5:5861. doi:10.1038/ncomms6861.

52. Li H, Durbin R. Fast and accurate short read alignment with Burrows-Wheeler transform. Bioinformatics. 2009;25(14):1754–1760. doi:10.1093/bioinformatics/btp324.

53. DePristo MA, Banks E, Poplin R, Garimella KV, Maguire JR, Hartl C, et al. A framework for variation discovery and genotyping using next-generation DNA sequencing data. Nature genetics. 2011;43(5):491–498. doi:10.1038/ng.806.

54. Yang J, Benyamin B, McEvoy BP, Gordon S, Henders AK, Nyholt DR, et al. Common SNPs explain a large proportion of the heritability for human height. Nature genetics. 2010;42(7):565. doi:10.1038/ng.608.

55. Yang J, Zaitlen NA, Goddard ME, Visscher PM, Price AL. Advantages and pitfalls in the application of mixed-model association methods. Nature genetics. 2014;46(2):100. doi:10.1038/ng.2876.

56. Kim D, Pertea G, Trapnell C, Pimentel H, Kelley R, Salzberg SL. TopHat2: accurate alignment of transcriptomes in the presence of insertions, deletions and gene fusions. Genome biology. 2013;14(4):1. doi:10.1186/gb-2013-14-4-r36.

57. Wagner GP, Kin K, Lynch VJ. Measurement of mRNA abundance using RNA-seq data: RPKM measure is inconsistent among samples. Theory in Biosciences. 2012;131(4):281–285. doi:10.1007/s12064-012-0162-3.

58. Ramaswami G, Lin W, Piskol R, Tan MH, Davis C, Li JB. Accurate identification of human Alu and non-Alu RNA editing sites. Nature methods. 2012;9(6):579. doi:10.1038/NMETH.1982.

59. Browning BL, Browning SR. Genotype imputation with millions of reference samples. The American Journal of Human Genetics. 2016;98(1):116–126. doi:10.1016/j.ajhg.2015.11.020.

60. Panagiotou OA, Ioannidis JP, Project G4S. What should the genome-wide significance threshold be? Empirical replication of borderline genetic associations. International journal of epidemiology. 2011;41(1):273–286. doi:10.1093/ije/dyr178.

